# Conserved T-cell epitopes predicted by bioinformatics in SARS-COV-2 variants

**DOI:** 10.1101/2021.08.12.456182

**Authors:** Feiyu Lu, Shengnan Wang, Ying Wang, Yunpeng Yao, Yangeng Wang, Shujun Liu, Yangyang Wang, Yongli Yu, Liying Wang

## Abstract

**Background:** Finding conservative T cell epitopes in the proteome of numerous variants of SARS-COV-2 is required to develop T cell activating SARS-COV-2 capable of inducing T cell responses against SARS-COV-2 variants.

**Methods:** A computational workflow was performed to find HLA restricted CD8^+^ and CD4^+^ T cell epitopes among conserved amino acid sequences across the proteome of 474727 SARS-CoV-2 strains.

**Results:** A batch of covserved regions in the amino acid sequences were found in the proteome of the SARS-COV-2 strains. 2852 and 847 peptides were predicted to have high binding affinity to distint HLA class I and class II molecules. Among them, 1456 and 484 peptides are antigenic. 392 and 111 of the antigenic peptides were found in the conseved amino acid sequences. Among the antigenic-conserved peptides, 6 CD8^+^ T cell epitopes and 7 CD4^+^ T cell epitopes were identifed. The T cell epitopes could be presented to T cells by high-affinity HLA molecules which are encoded by the HLA alleles with high population coverage.

**Conclusions:** The T cell epitopes are conservative, antigenic and HLA presentable, and could be constructed into SARS-COV-2 vaccines for inducing protective T cell immunity against SARS-COV-2 and their variants.

## Background

Severe acute respiratory syndrome-coronavirus 2 (SARS-COV-2) causing Coronavirus disease 19 (COVID-19) (1, 2) has been epidemic in the world for more than 19 months, with more than 191 million people infected and more than 4 million death, reported by World Health Organization (WHO) on June 14, 2021. To fight this, a variety vaccines against SARS-COV-2 have been developed at unprecedented speed. Among them, 108 candidate vaccines are in clinical phase and 22 vaccines have been authorized for emergency use(3, 4). Noticeably, these SARS-COV-2 vaccines mainly target the spike (S) protein which binds the receptor angiotensin-converting enzyme 2 (ACE2) of host cells for the SARS-COV-2 entry, and are believed to be able to induce the neutralizing antibodies specific to the S protein, thereby providing immune protection in the individuals who received the SARS-COV-2 vaccines (5, 6). However, variants of SARS-CoV-2 with S protein mutations have emerged around the world. Up to August 3^rd^, 2021, 1,009,273 varied SARS-COV-2 genomes had been deposited into National Center for Biotechnology Information (NCBI) Virus Database (7). Among them, B.1.1.7,B.1.351, and P.1 were reported to be dominantly transmitted in the UK, South Africa and Brazil, respectively, and have spread to several other countries. The SARS-CoV-2 variants evolve as a result of mutation and natural selection for their favorable traits, one of which is to evade the host immunity, such as neutalizing antibodies, induced by infection or vaccines, raising concern on a growing number of reinfection and a reduction in the effectiveness of SARS-CoV-2 vaccines in use (8–10). Evidently, the sporadic reinfection of SARS-COV-2 reported in USA, Belgium, Hong Kong of China and Ecuador (11–15) could be correlated to the SARS-COV-2 variants because of the varied genomes of SARS-COV-2 from first and second episodes of the infections. Recent studies revealed that the B.1.351 variant dodged the neutralizing antibodies induced by several SARS-COV-2 vaccines, indicating that the mutations could change the B cell epitopes in the S proteins (15–17) and thus calling for upgrading the SARS-COV-2 vaccines against the established and emerging variants.

Recently, vaccine-induced T cell responses have been noticed for up-grading SARS-COV-2 vaccines for resistant variants. Clinical studies showed that some cases of asymptomatic SARS-COV-2 exposure have been associated with cellular immune response without seroconversion, indicating that SARS-CoV-2 specific T cells could be relevant in disease control even in the absence of neutralizing antibodies. In COVID-19 patients, neutralizing antibody titers were not generally paralleled with reduced disease severity, but SARS-CoV-2 specific CD4^+^ and CD8^+^ T cells are each associated with milder disease (18–20). The data suggest that immunological memory of T cells, if efficiently induced by the SARS-COV-2 vaccines, could provide protection. Intrestingly, unlike B cells, which mainly target accessible proteins like S proteins or nucleocapsid phosphoprotein (N proteins) in SARS-COV-2, T cells can target all of viral proteins in SARS-COV-2 proteome. More importantly, the T cell responses could be less affected by the mutations, and some of T cell epitopes distributed in the SARS-COV-2 proteome seem more stable than the B cell epitopes (21, 22). Therefore, SARS-CoV-2 vaccines capable of activating both CD4^+^ and CD8^+^ T cells are likely to induce the protection of SARS-CoV-2 variants. If so, it is required to develop the T cell vaccines, targeting the mutation-resistant T cell epitopes in the SARS-CoV-2 proteome.

Here, we systemically compare and analyze T cell epitopes in the initially identified SARS-CoV-2 and in the rapidly accumulated SARS-CoV-2 variants, with the aim of locating some of consevative T cell epitopes in the SARS-CoV-2 proteome for developing T cell vaccines to fight the SARS-CoV-2 variants.

## Methods

### Data retrieval

The complete sequence of SARS-CoV-2 isolate Wuhan-Hu-1 was retrieved from the nucleotide database available at NCBI with the accession number NC_045512.2 (23). It encodes 9860 amino acids translating into several non-structural proteins (NSP) like NSP1-16 and accessory proteins like open reading frame (ORF) 3a, 6, 7a, 7b, 8, 10, along with structural proteins including S protein, envelope (E) protein, membrane glycoprotein (M protein), and N protein. The accession number in the NCBI of these proteins are shown in Table S1.

### Selection of human leukocyte antigen (HLA) class I and HLA class II alleles

HLA class I and II alleles were selected on the basis of their occurrence worldwide. The worldwidely most frequent 27 HLA class I and 26 HLA class II alleles (24, 25), which accouting for the population coverage > 97% and >99% worldwide, rspectively. As shown in Table S2.

### Alignments of SARS-COV-2 strains

474727 strains of SARS-CoV-2 sequences depositing to the Global Initiative of Sharing All Influenza Data (GISAID) database (26, 27) with the presenting date ranging from January 27^th^, 2021 to April 27^th^, 2021 on the date of April 27^th^ 2021, were collected. Using the server of COVID-19 CoV Genetics (COVID CG) (27), the amino acid sequences of each protein of repertoire to the corrsponding amino acid suquences of hCoV-19/Wuhan/WIV04/2019 (MN996528.1), which sharing 100% homology with the reference sequence NC_045512.2 (23) were aligned. High frequency variations were defined as the ratio of occuring counts of amino acid mutations to 474727 greater than 10^−3^ (28).

### Prediction of SARS-CoV-2-derived CD8^+^ T cell and CD4^+^ T cell epitopes

The sequences of 26 proteins encoded by the SARS-COV-2 genome were split into 9 amino acid-long peptides for CD8^+^ T cell epitopes and 15 amino acid-long peptides for CD4^+^ T cell epitopes covering the complete proteome of the SARS-COV-2. The NetMHCpan 4.1(29) server was utilized to screen the 9-mer peptides across the proteome of the SARS-COV-2 (NC_045512.2) for their binding affinity with distinct HLA class I molecules encoded by selected 27 HLA class I alleles. The prediction of binding affinity was based on more than 850,000 quantitative binding affinity and mass-spectrometry eluted ligands peptides. The NetMHCpan 4.1 server analysis resulted in the evaluation about the binding affinity and the binding strength between 9-mer peptides and selected HLA class I molecules, expressed by cores of nanomolar IC50 and percentile rank respectively. Thresholds for high binding affinity were set ≤500 nM and strong binding strength were set at top 0.5%. The NetMHCIIPan 4.0 server (29) was used to predict the binding affinity and binding strength of the peptides (15 mer) across the proteome of the SARS-COV-2 (NC_045512.2) to the selected HLA class II molecules (Table S2). Thresholds for high binding affinity were set ≤500 nM and strong binding strength were set at top 1%. The antigenicity of screened peptides to both HLA class I and class II molecules was predicted using Vaxijen-2.0 (30, 31), which analyzed higher order interactions positions of protein sequence and exploiting the physicochemical properties (hydrophobicity, molecular size, polarity) of amino acids. Each of the peptides was scored for selecting. The peptides with the scores above 0.4 were determined antigenecity. Exclude the peptides located at the mutated positions based on the alignment of 474727 strains depositted to the GISAID database. The peptides presented by more HLA alleles (≥3 alleles) were rescreened with the hypothesis that increased HLA binding promiscuity meant broader population (32). The fraction of individuals capable of responding to each of the candidate T cell epitopes was calculated by IEDB population Coverage (33) based on the coresponding HLA genotypic frequencies.

### Molecular docking of HLA molecules to CD8^+^ T cell epitopes and CD4^+^ T cell epitopes

The spacial structure of HLA molecules were downloaded from pHLA database (34). Docking simulations between predicted CD8^+^ T cell epitopes or CD4^+^ T cell epitopes and presented HLA molecules were performed using the GalaxyPepDock server (35), which enables prediction of 3D protein-peptide complex structure interactions from input protein structure and peptide sequence using similar interactions found in the structure database and energy-based optimization. The approaches of graphing are python and Graphpad.

## Results

### Alignment of sequences of proteins encoded by SARS-COV-2 strains

To develop T cell vaccines, targeting the mutation-resistant T cell epitopes in the SARS-CoV-2 proteome, we tried to screen universal T cell epitopes in the coserved amino acid sequences across SARS-COV-2 proteome. Firstly, we collected and comprehensively screened all of the 474727 SARS-COV-2 sequences with the presenting date ranging from January 27^th^, 2021 to April 27^th^,2021, deposited to GISAID database (26, 27). These strains were isolated from the Asia (9403/474727), Africa (961/474727), Europe (315833/474727), Oceania (1225/474727), North America (144760/474727) and South America (2545/474727). To find the conserved amino acid sequences in the repertoire of the SARS-COV-2, using the server of COVID CG (27), we aligned the proteome of all these submitted sequences with the sequence of hCoV-19/Wuhan/WIV04/2019 (MN996528.1) (36) which shares 100% homology with the reference sequence NC_045512.2 (23). The alignment revealved that the amino acid mutations were widely distributed in the proteins across the SARS-COV-2 proteome, ranging from 2.12×10^−6^ to 9.96×10^−1^ in occuring frequency. The frequency higher than 10^−3^ was defined as relatively high frequency mutations (28). With this cut-off, we identified 899 amino acid substitutions and 20 amino acid deletions as relatively high frequency variations at 816 positions in the SARS-COV-2 proteome (Figure 1A). As for the positions, 117 are in the S protein, a target for inducing neutralizing antibodies; 153 in NSP3; 66 in NSP6 and ORF3a; 53 in NSP2 and 1 to 42 in the other proteins encoded by the SARS-COV-2 genome. Howerver, neverthless, we identifed a batch of positions which are covserved in the proteome of SARS-COV-2 (Figure 1B). Specifically, 1156 are in the S protein; 1792 and 903 in the NSP3 and NSP12 proteins, 12 to 585 in the rest proteins of SARS-COV-2 proteome.

**Figure 1.**
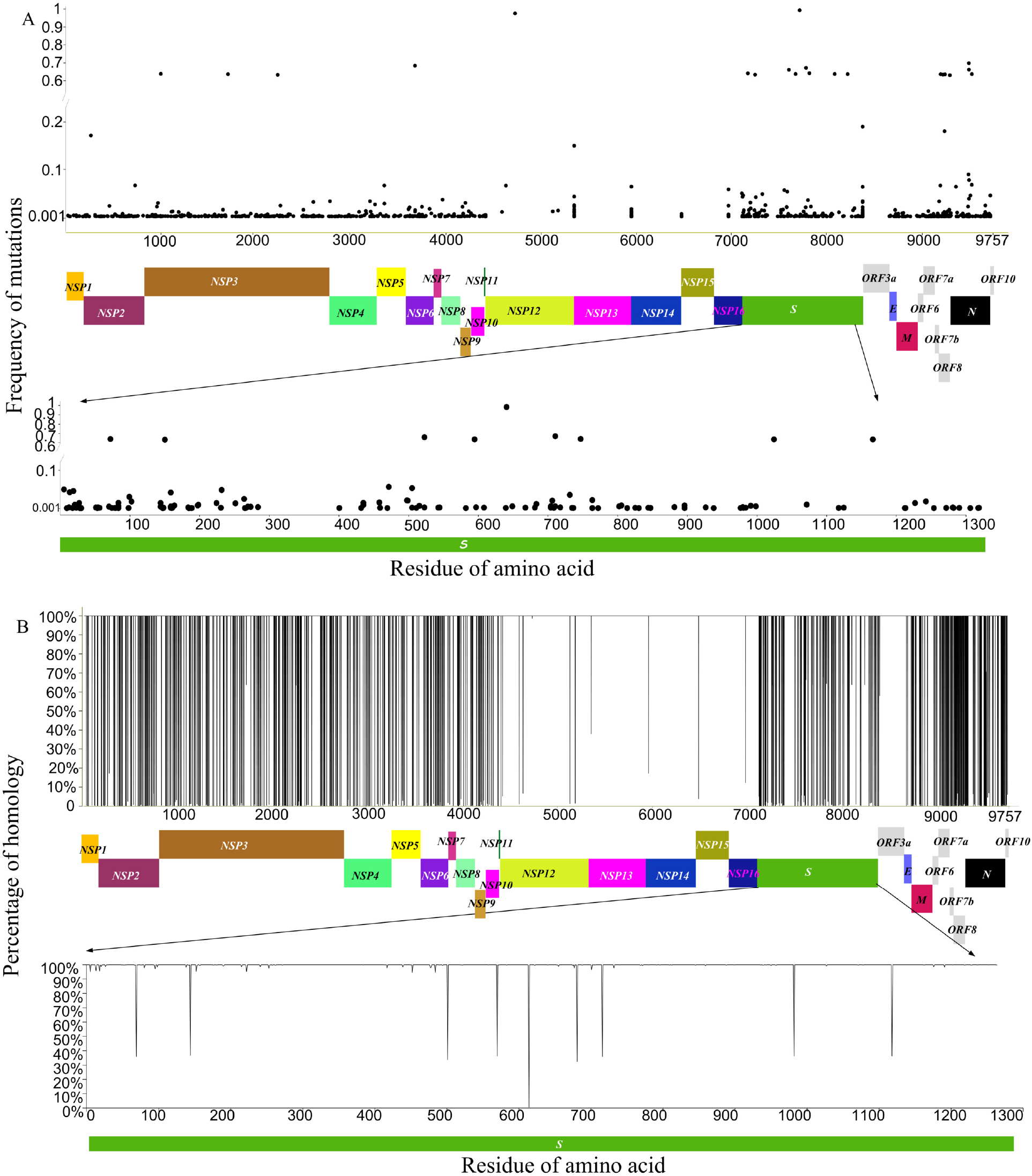
Mutated and conserved amino acid sequences of SARS-COV-2. Frequency of mutation and homology plot based on the full-length proteome sequence of SARS-CoV-2. Proteome sequences of 474727 SARS-CoV-2 strains deposited to the GISAID database with the presenting date ranging from January 27th, 2021 to April 27th, 2021 were aligned with hCoV-19/Wuhan/WIV04/2019 (accession number MN996528.1), which shares 100% homology with the reference sequence Wuhan-Hu-1(NC_045512.2). Mutations with occurring frequency greater than 10^−3^ were recruited. 899 amino acid substitutions and 20 amino acid deletions were found across the proteome of SARS-COV-2. (**A**) Dots represent the amino acid of mutations. (**B**) Lines represent the amino acid of mutations. The percentage of homology 100% represent the conserved amino acid sequences. S (spike protein), E (envelope protein), M (membrane glycoprotein), N (nucleocapsid phosphoprotein). NSP (nonstructrural protein). ORF (open reading frame).

### Prediction of CD8^+^ T cell and CD4^+^ T cell epitopes in the conserved amino acid sequences across SARS-CoV-2 proteome

Next, we screened and predicted T cell epitopes in the repertoire of the SARS-COV-2 proteome. As for the CD8^+^ T cell epitopes, using NetMHCpan 4.1 server (29), we screened the 9-mer peptides across the proteome of the SARS-COV-2 (NC_045512.2) for their binding affinity with distinct HLA class I molecules encoded by 27 HLA class I alleles (Table S2). The alleles account for the population coverage > 97% worldwide (24). Based on more than 850,000 quantitative binding affinity and mass-spectrometry eluted ligands peptides, the NetMHCpan 4.1 server analysis resulted in the evaluation about the binding affinity and the binding strength between 9-mer peptides and selected HLA class I molecules, expressed by scores of nanomolar IC50 and percentile rank respectively. Thresholds for high binding affinity were set ≤500 nM and strong binding strength were set at top 0.5%. Eventually, 2852 of the peptides with high affinity to the distinct class I molecule were selected for further analysis. As for the CD4^+^ T cell epitopes, using the NetMHCIIPan 4.0 server (29), we predicted the binding affinity and binding strength of the peptides (15 mer) across the proteome of the SARS-COV-2 (NC_045512.2) to the HLA class II molecules (Table S2) encoded by the elleles which account for the population coverage >99% (25). Resultantly, 847 candidate HLA class II binding peptides from the SARS-COV-2 proteome were predicted to have high binding affinity (≤ 500 nM) and strong binding strength (percentile rank ≤1%).

Furthermore, we predicted wheather these candidate peptides paired with distinct HLA molecules could efficiently bind T cell recepters (TCR) by analysing hydrophobicity, molecular size, polarity of amino acids of the pepetids, using Vaxijen-2.0 (30, 31). Each of the peptides was scored by the server. The peptides scored above 0.4 were selected as the antigenic ones which turned out to be 1456 pepetides and 484 of pepetides, restricted by the selected HLA class I and class II molecules, respectively. Based on the above screening, we tried to locate T cell epitopes in the conserved amino acid sequences across the proteome of SARS-COV-2 by aligning the sequences of the selected T cell epitopes with the SARS-COV-2 sequences deposited to the GISAID database from January 27th, 2021 to April 27th, 2021. As shown in the Figure 2A, 392 T cell epitopes restricted to the selected HLA class I molecules were found. Specifically, 82 are in the NSP3, 69 in the NSP12, 43 in the S protein, 30 in the NSP13, 21 in the NSP4, 20 in the M protein,18 in the NSP15, 17 in the NSP2 and NSP16, 14,13 and 11 in the NSP14, NSP6 and N protein, 8 and 7 in the NSP5 and NSP8, 1 to 5 are in the ORF6, NSP10, ORF3a, NSP7, E protein, NSP1 and NSP9. And 111 candidate T cell epitopes restricted to selected HLA class II molecules were found. Among them, 34 are in the NSP3, 16 in the S protein, 15 in the NSP12, 13 in the NSP13, 11 in the NSP15, 1 to 4 in the NSP1-2,NSP4-8, NSP10, NSP14, NSP16, N and M proteins.

**Figure 2.**
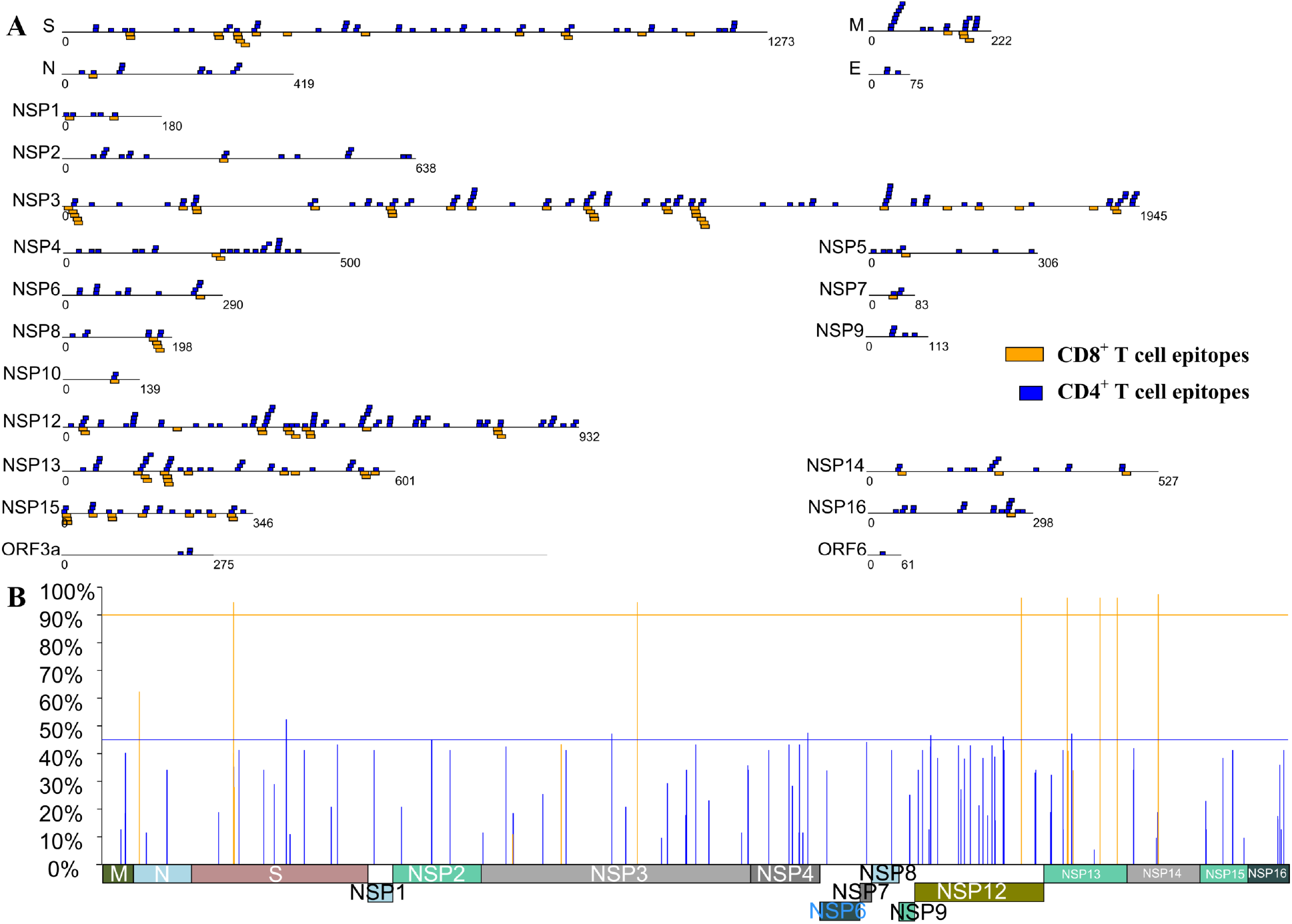
The predicted conserved and universal T cell epitopes. 9-mer peptides (in blue) derived from SARS-CoV-2 (NC_045512.2) presented by selected HLA class I molecules with high binding affinity (IC50 ≤ 500 nM, percentile rank < 0.5%) and antigenicity predicted by the NetMHCpan 4.1 and Vaxijen-2.0 server. 15-mer Peptides (in yellow) across the proteome of the SARS-COV-2 (NC_045512.2) presented by the selected HLA class II molecules with high binding affinity (IC50≤500 nM, percentile rank < 1%) and antigenicity predicted by the NetMHCIIPan 4.0 and Vaxijen-2.0 server. (**A**) Squares represent the predicted peptides located at the conserved amino acid sequences.(**B**) The vertical lines represent the peptides loacting at conserved amino acid sequences presented by more alleles of HLA molecules (≥3 alleles). The population coverage was calculated by IEDB population Coverage server. Transverse lines represent the cut-off of population coverage 45% or 90%, respectively. S (spike protein), E (envelope protein), M (membrane glycoprotein), N (nucleocapsid phosphoprotein). NSP (nonstructrural protein). ORF (open reading frame).

To find the universal T cell epitopes, we further assessed the HLA binding promiscuity of the selected T cell epitopes. Firstly, the T cell epitopes presented by one or two HLA molecules were excluded. As shown in the Figure 2B, 88 or 14 T cell epitopes restricted by more than two HLA class I or II molecules were selected as the candidate CD8^+^ T cell epitopes or CD4^+^ T cell epitopes, respectively. Then, based on the coresponding HLA genotypic frequencies, the fraction of individuals capable of responding to each of the candidate T cell epitopes was calculated by IEDB population Coverage (33). The results showed that the potential coverage in populations was ranging from 5.38% to 52.34% for the 88 CD8^+^ T cell epitopes, and 27.96% to 97.48% for 14 CD4^+^ T cell epitopes (Figure 2B). 6 of CD8+ and 7 of CD4+ epitopes were selected finally, with the population coverage > 45% for CD8^+^ T cell epitopes and > 90% for CD4^+^ T cell epitopes respectively.

### The interactions between predicted T cell epitopes and presented HLA molecules

Ultimately, we identified 6 CD8^+^ T cell epitopes, designated as S 691-699, NSP3 950-958, NSP4 420-428, NSP12 123-131, NSP12 647-655 and NSP13 209-217, and 7 CD4^+^ T cell epitopes, designated as S 310-324, NSP3 1134-1148, NSP12 778-792, NSP13 177-191, NSP13 413-427, NSP13 538-552, and NSP14 232-246, respectively. Specifically, among the CD8^+^ T cell epitopes, 4, 3 or 2 can be presented by the HLA class I proteins encoded by the alleles of HLA A*02:01, A*02:03 and A*32:01, of HLA A*02:06 and B*08:01, or of HLA A*68:02; A*03:01, A*11:01, A*30:02, B*15:01 and B*35:01 (Figure 3A), respectively. All of the CD4^+^ T cell epitopes can be presented by the HLA class II proteins encoded by the alleles of HLA DPA10103-DPB10401, DPA10201-DPB10101 and DPA10301-DPB10402. Also, among the CD4^+^ T cell epitopes, 5 or 1 can be presented by HLA class II proteins encoded by the alleles of DPA10103-DPB10201 or DPA10201-DPB10501, respectively (Figure 3B).

**Figure 3.**
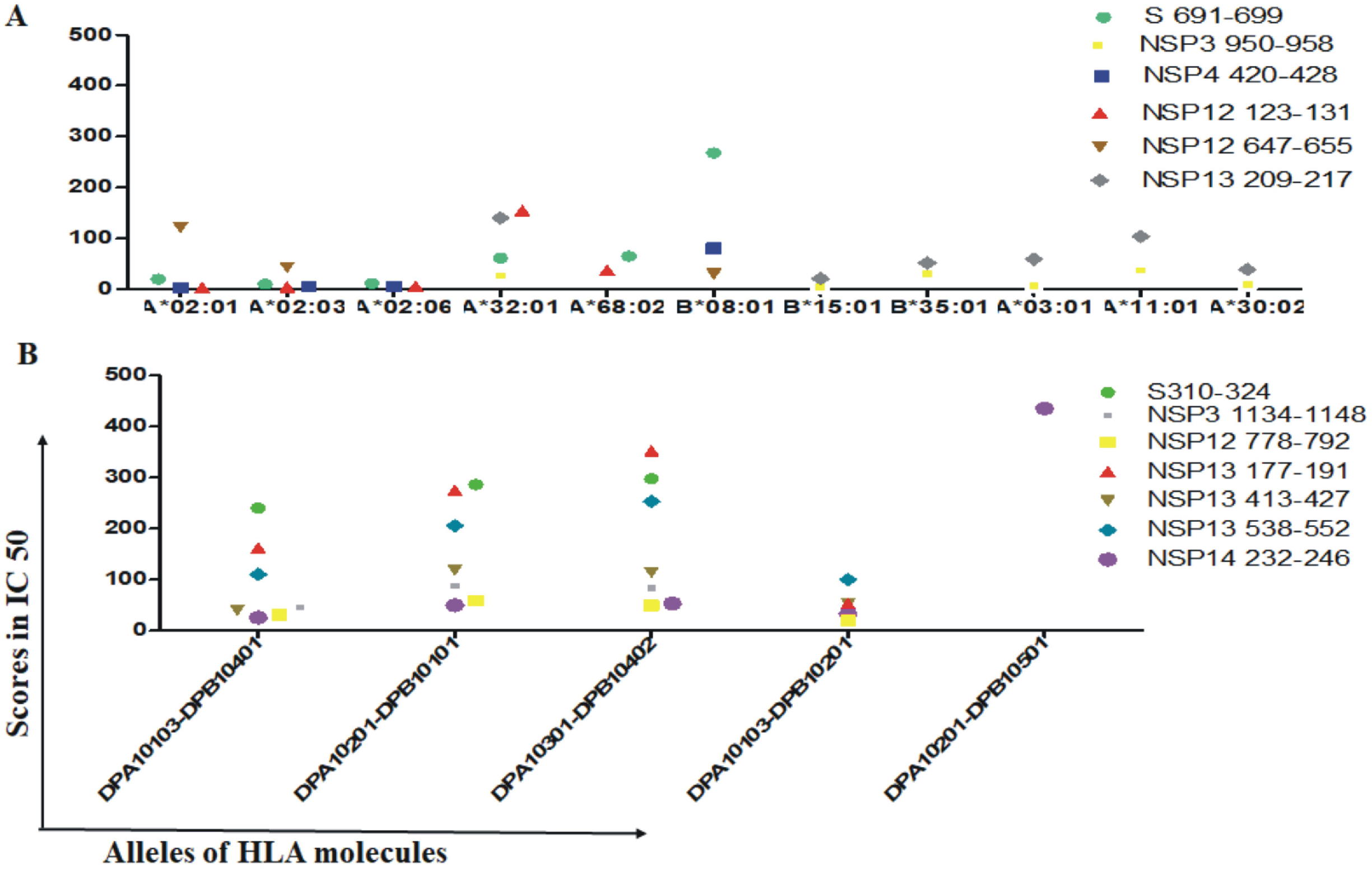
The selected conservative and universal T cell epitopes and the presented HLA molecules. The selected conservative and universal T cell epitopes with antigenicity and high binding affinity to presented HLA molecules illustrated by dots. (**A**) CD8^+^ T cell epitopes. (**B**) CD4^+^ T cell epitopes.

At last, we conducted the docking simulations between the selected T cell epitopes and the HLA molecules using the GalaxyPepDock (35), a server which enables prediction of 3D protein-peptide complex structure interactions from input protein structure and peptide sequence using similar interactions found in the structure database and energy-based optimization. Ten models of each epitope-HLA complex were generated on the basis of minimized energy scores. The scores of estimated accuracy represent the estimated fraction of correctly predicted binding site residues. We docked each of the selected CD8^+^ T cell epitopes or CD4^+^ T cell epitopes with the corresponding HLA molecules and scored the pairs. As shown in Table 1, all of the pairs between the 6 selected CD8^+^ T cell epitopes and the distinct HLA class I molecules scored with estimated accuracy 1. The pairs between the 7 selected CD4^+^ T cell epitopes and the distinct HLA class II molecules scored with estimated accuracy in a range from 0.792 to 0.852, which presents high grade 3D fits of the pairs. The results were further supporting that these epitopes should be strong binders to presented HLA molecules and promising candidates for vaccine development studies.

**Table 1.**
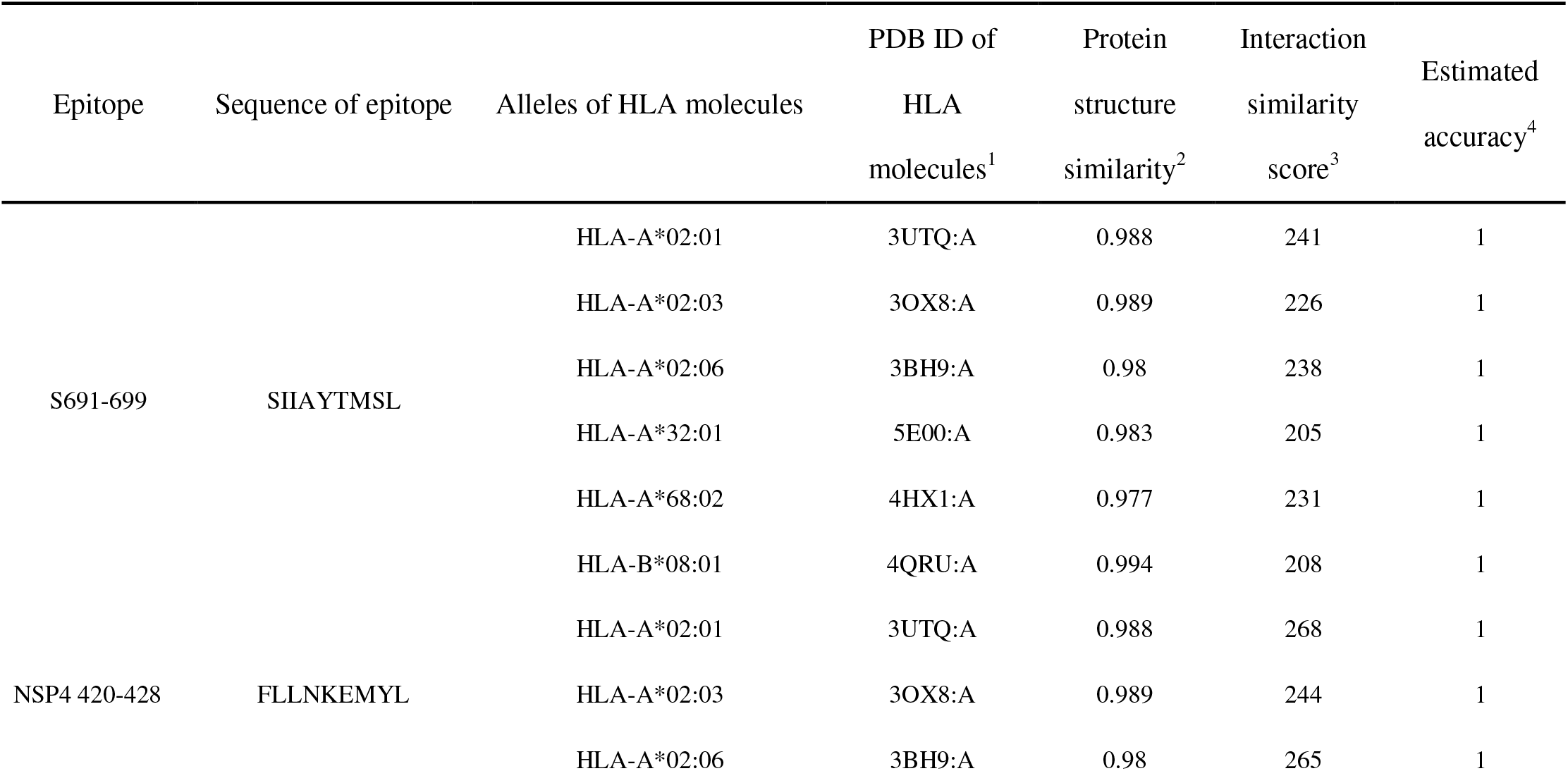
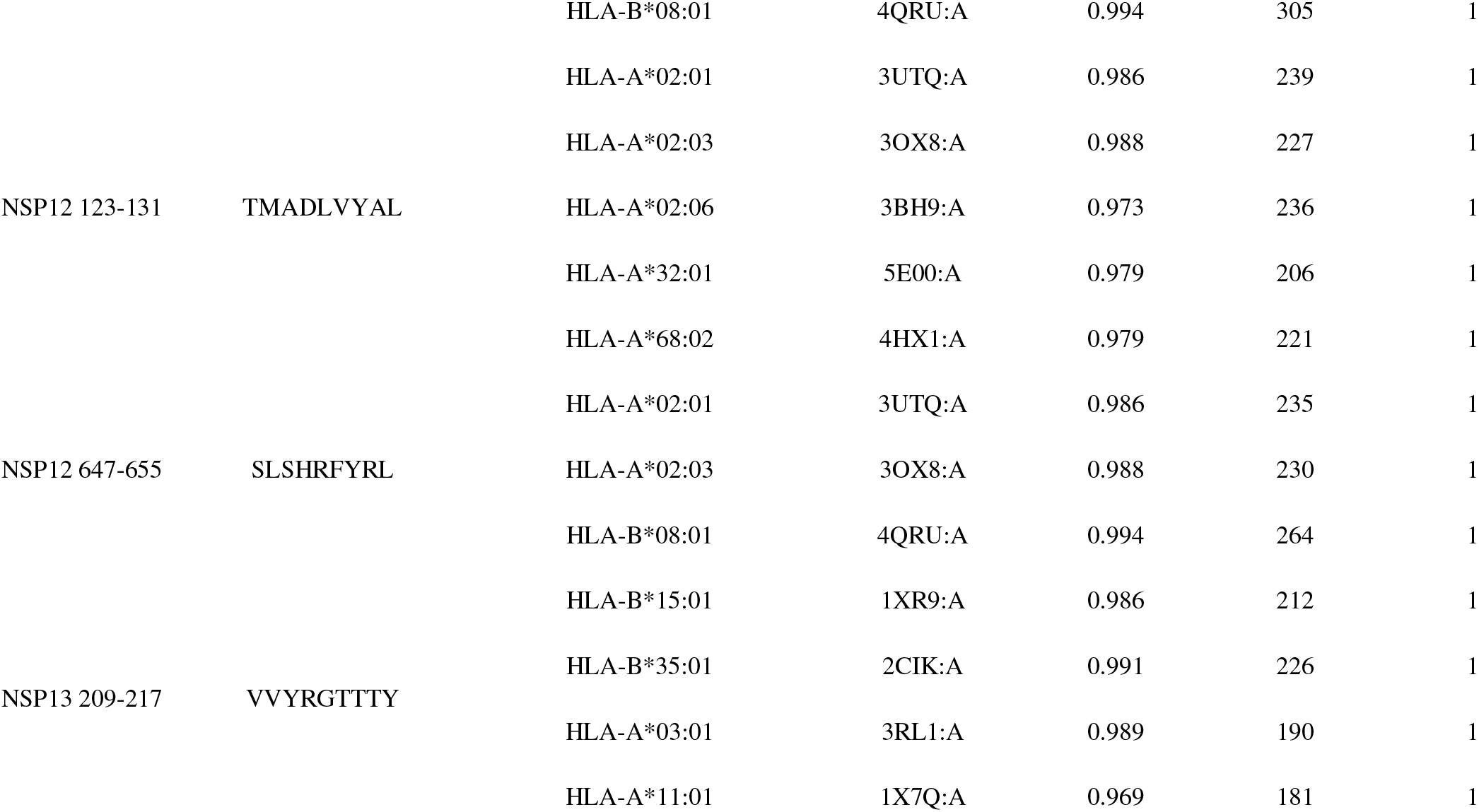
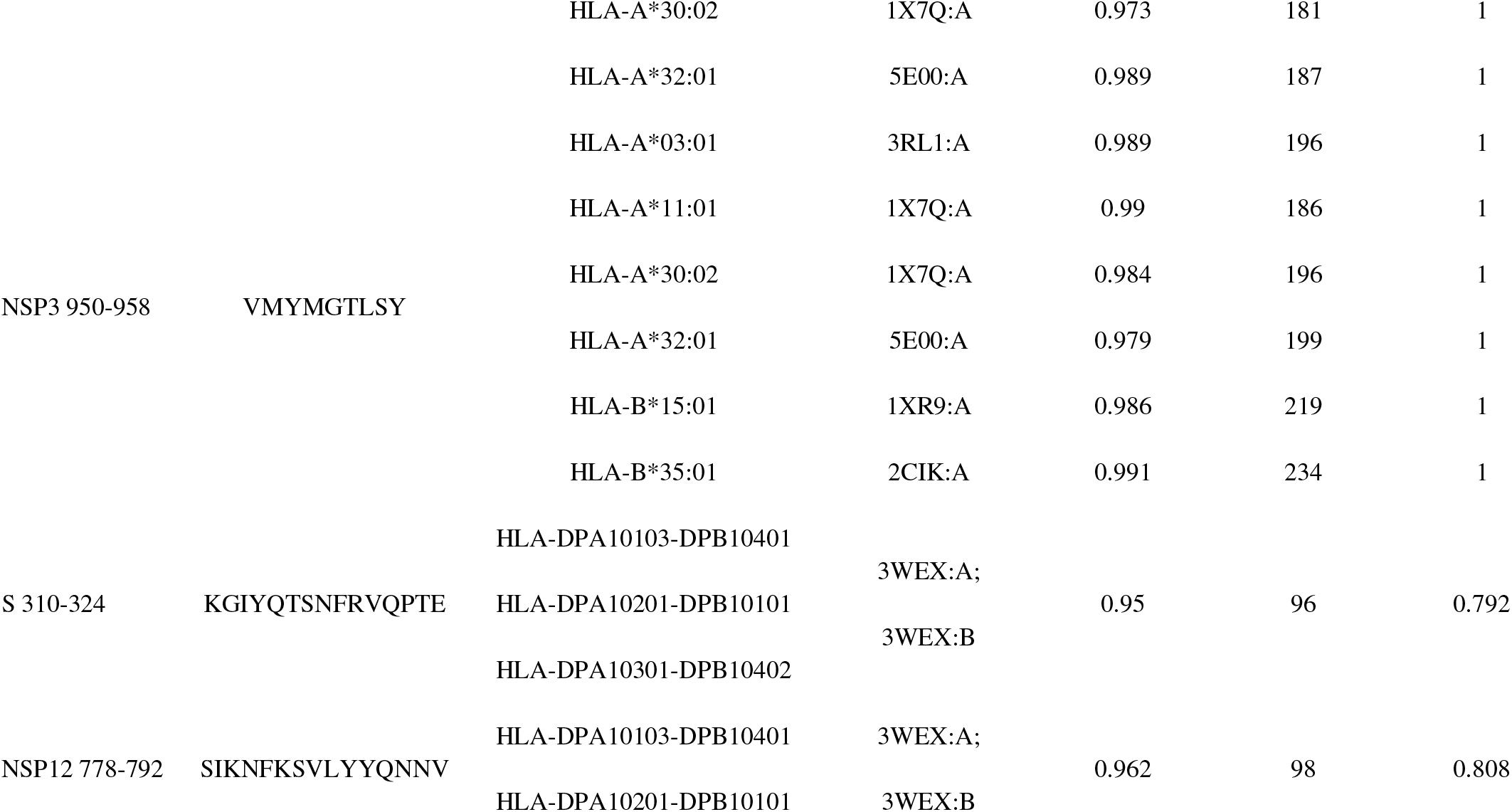
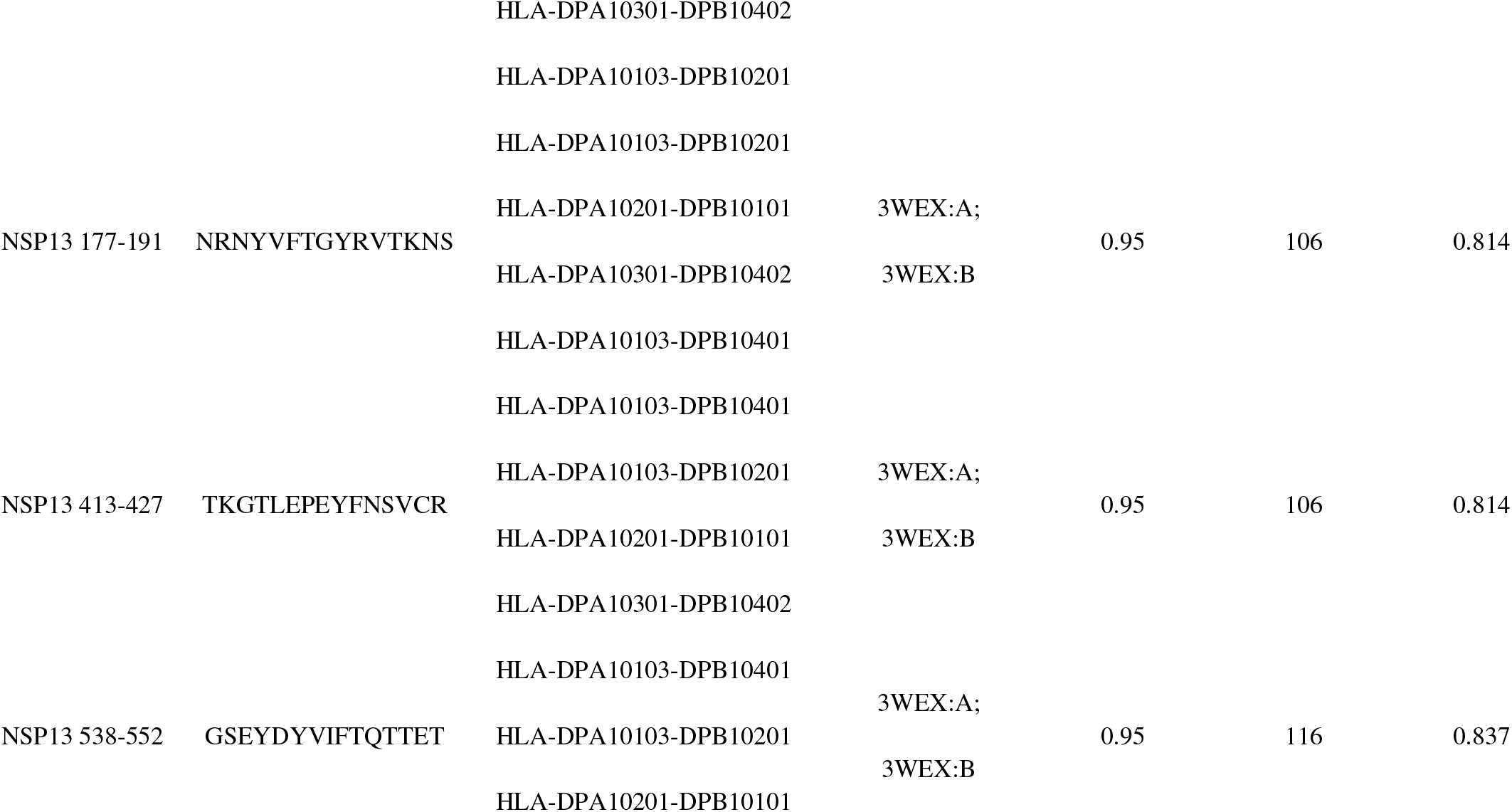
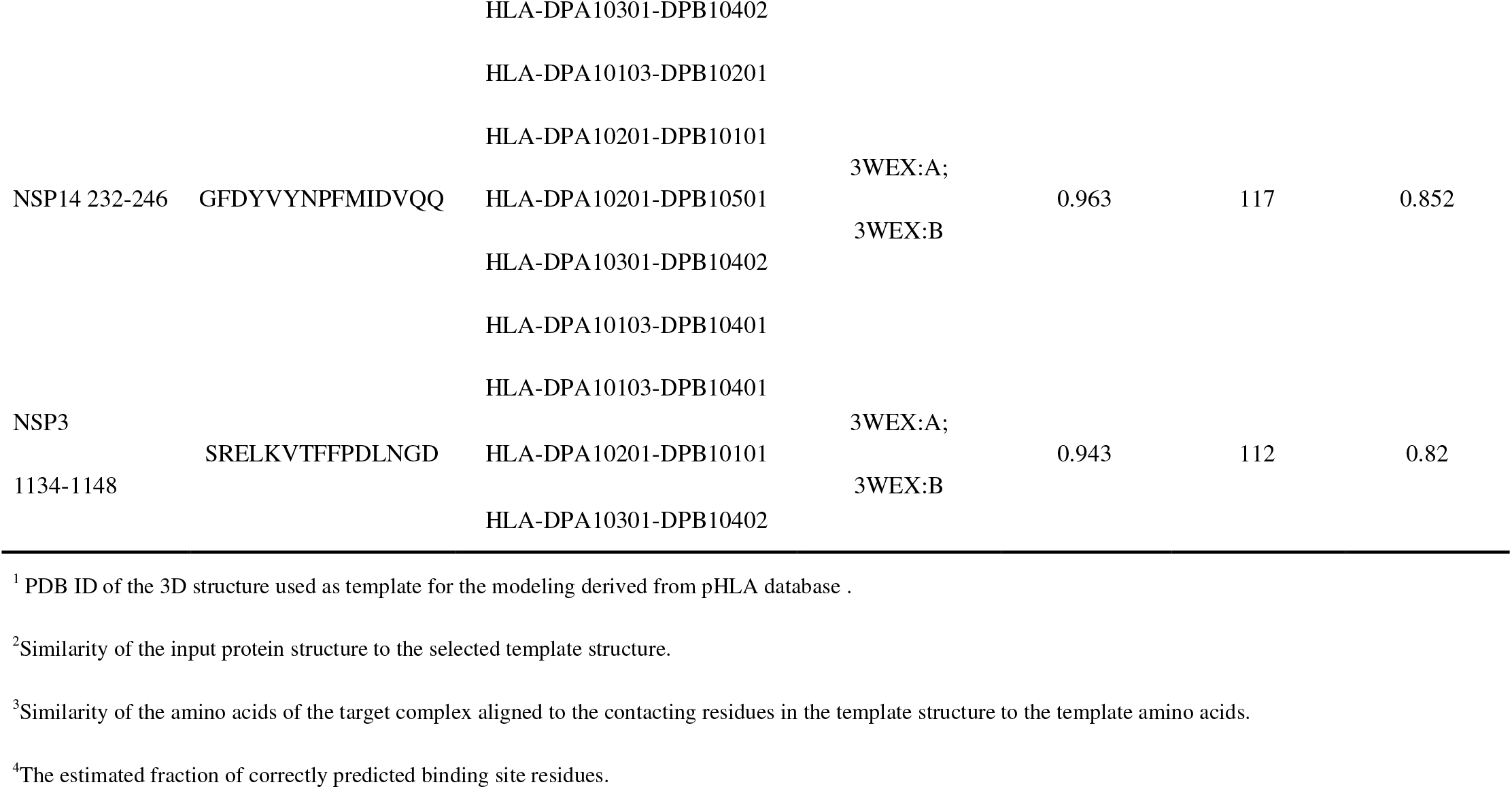
Docking simulations between the selected T cell epitopes and the presented HLA molecules by GalaxyPepDock.

## Discussion

Using the bioinformatic tools, we screened and identified 6 CD8^+^ T cell epitopes and 7 CD4^+^ T cell epitopes in the conserved amino acid sequences among the 474727 SARS-COV-2 strains whose sequences are deposited to the GISAID. Notably, the inter-individual variation in the SARS-CoV-2 sequences is low, compared to many other RNA viruses (37, 38), in part because coronaviruses encode a 3’–5’ exonuclease activity (NSP14), which provides a proofreading function that enhances replication fidelity and limits viral sequence diversification (39). Thus, it is virtually possible to find T cell epitopes in the repertoire of the SARS-COV-2 proteins. Primarily, these T cell epitopes exist at the the essential amino acid sequences which are crutial for the SARS-CoV-2 life-cycle in the host cell. As shown in Fig 2B, two conserved T cell epitopes were identifed in S proteins among the 474727 SARS-CoV-2 strains. One is a CD4^+^ T cell epitope (S310-324) which is partially overlapped with the receptor binding domain (RBD) of the S1 subunit (S319-541), and another is a CD8^+^ T cell epitope (S691-699) which locates in the S1/S2 cleavage region (S690-709). The SARS-CoV-2 with mutations in the epitopes (S310-324, S691-699) may hamper the cell entry of the SARS-CoV-2. Two T cell epitopes were identified in the NSP3, a papain-like protease which is critical for the SARS-CoV-2 to yield mature functional proteins from a polyprotein (40, 41). One is CD8^+^ T cell epitope (NSP3 950-958) which locates in the crutial domain of the papain-like protease. Another is CD4^+^ T cell epitope (NSP3 1134-1148) which locates in a nucleic acid-binding domain (NAR). The SARS-CoV-2 with mutations in the epitopes (NSP3 950-958, NSP3 1134-1148) may lose its capacity to generate functional proteins. Two CD8^+^ T cell epitops (NSP12 123-131; NSP12 647-655) and one CD4^+^ T cell epitope (NSP12 778-792) are in the NSP12 protein, a RNA-dependent RNA polymerase (RdRp) which is required for generating SARS-CoV-2 RNAs (42). Three CD4^+^ T cell epitopes (NSP13 177-191; NSP13 413-427; NSP13 538-552) and one CD8^+^ T cell epitope (NSP13 209-217) is in the NSP13 protein, a helicase which can unwind DNA or RNA in an nucleotide triphosphate (NTP)-dependent manner with a 5’>3’ polarity (43), a process critical for SARS-CoV-2 replication-transcription. One CD4^+^ T cell epitope (NSP14 232-246) is in the NSP14 protein, a 3’–5’ exonuclease which provids a proofreading function that enhances replication fidelity and limits viral sequence diversification (44); One CD8^+^ T cell epitope (NSP4 420-428) is identified in the NSP4 protein, a membrane-spanning protein containing transmembrane domain 2 which helps the SARS-CoV-2 to modify endoplasmic reticulum (ER) membrane in the host cell (45). Obviously, the 6 CD 8^+^ T cell epitopes and 7 CD4^+^ T cell epitopes identified in this study are all located in the key proteins for SARS-CoV-2 life-cycle and therefore could be the universial T cell epotopes which are conservatively retained in all SARS-CoV-2 strains.

Hopefully, the conserved T cell epitopes presented here could be constructed into SARS-CoV-2 vaccines to induce more univerisal T cell responses aginst the SARS-CoV-2. Accumulating evidence showed that the T cell immunity, if being induced by the SARS-CoV-2 vaccines contained the T cell epitopes, could be benefitial to the patients infected with SARS-CoV-2. As reported, circulating SARS-CoV-2-specific CD8^+^ T cells and CD4^+^ T cells were identified in ~70% and 100% of COVID-19 convalescent patients, they are associated with the better outcomes of the COVID-19 patients (18, 46). Importantly, the protective specific T cell responses against the SARS-CoV-2, after being induced, could be more sustained compared to the neutralizing humoral responses (47). Functional SARS-CoV-2-specific T cell responses are retained at 6 months following infection (48). In contrast, SARS-CoV-2-specific antibody responses were waned after 1 month after symptom onset (49). Furthermore, SARS-CoV-2-specific memory CD4^+^ T cells, if being induced prior to the natural infection, will help the SARS-CoV-2-specific B cell to launch rapid and robust antibody responses (46). Intrestingly, the SARS-CoV-2 needs to maintain the T cell epitopes revealed in this study for its life-cycle, even facing the evolution pressures caused by the acquired specific immunity induced by the natural infection of vaccination. Thus, the SARS-CoV-2 vaccines constructed with the T cell epitopes identified in this study could induce more univerisal T cell responses aginst the SARS-CoV-2 variants. Lastly, the revealed T cell eptitopes, due to their capacity of being presented by the HLA molecules encoded by widely distributed HLA alleles, could be ideal targets for developing novel SARS-CoV-2 vaccines which induce the protective T cell immunity in large populations worldwide.

## Acknowledgments

The authors would like to thank Zhengang Jiao for his help in the application of informatics software, as well as Mengyuan Kou, Wenting Lu, and Cuiyun Cui for their help to the authors in other aspects.

